# Subpopulation identification for single-cell RNA-sequencing data using functional data analysis

**DOI:** 10.1101/760413

**Authors:** Kyungmin Ahn, Hironobu Fujiwara

## Abstract

**Background:** In single-cell RNA-sequencing (scRNA-seq) data analysis, a number of statistical tools in multivariate data analysis (MDA) have been developed to help analyze the gene expression data. This MDA approach is typically focused on examining discrete genomic units of genes that ignores the dependency between the data components. In this paper, we propose a functional data analysis (FDA) approach on scRNA-seq data whereby we consider each cell as a single function. To avoid a large number of dropouts (zero or zero-closed values) and reduce the high dimensionality of the data, we first perform a principal component analysis (PCA) and assign PCs to be the amplitude of the function. Then we use the index of PCs directly from PCA for the phase components. This approach allows us to apply FDA clustering methods to scRNA-seq data analysis.

**Results:** To demonstrate the robustness of our method, we apply several existing FDA clustering algorithms to the gene expression data to improve the accuracy of the classification of the cell types against the conventional clustering methods in MDA. As a result, the FDA clustering algorithms achieve superior accuracy on simulated data as well as real data such as human and mouse scRNA-seq data.

**Conclusions:** This new statistical technique enhances the classification performance and ultimately improves the understanding of stochastic biological processes. This new framework provides an essentially different scRNA-seq data analytical approach, which can complement conventional MDA methods. It can be truly effective when current MDA methods cannot detect or uncover the hidden functional nature of the gene expression dynamics.

## Background

Single-cell RNA-sequencing (scRNA-seq) data analysis has been widely used to explore and measure the genome-wide expression profile of individual cells. Since the number of bioinformatics tools for scRNA-seq data analysis is growing dramatically, there are many studies comparing several statistical methods for scRNA-seq data analysis. Menon [35] reviewed three statistical clustering algorithms for scRNA-seq data to explicitly demonstrate their different behaviors in low- and high-read-depth data. Recently, Andrews and Hemberg [2] compared 12 clustering techniques on scRNA-seq data sets illustrating that the different methods generally produced clustering with minimal overlap. Duò et al. [11] extended these initial studies to 14 clustering algorithms on a total of 12 different simulated and real data sets showing the large differences in performance across data sets and clustering methods.

These statistical methods and algorithms for scRNA-seq data analysis belong to the general framework of *multivariate data analysis* (MDA), which helps analyze the gene expression data to understand stochastic biological processes. However, several shortcomings have arisen when the data are treated as vectors of discrete samples instead of continuous samples. One of the great advantages of applying this framework, i.e., *functional data analysis* (FDA) [47], is that we consider the *dependency* or *connectivity* between the samples. Several works are performed on gene expression data by applying FDA to increase the performance of statistical analysis [30, 10, 38, 29, 32, 3, 61, 60].

Based on these ideas, we propose an FDA technique for scRNA-seq data analysis to improve the accuracy of the classification of cell types. An important aspect of this study is that we view multivariate gene expression data as functional gene expression data. The different point of view from standard multivariate data analysis underlies the structure of raw observations being functional. This approach allows us to detect the functional nature of scRNA-seq data and uncover the functional characteristics of cell populations. This eventually classifies the subpopulations of cell types that cannot be detected by standard multivariate statistical methods. Given that a function does not allow the permutations of phase components of a function, we directly use principal components (PCs) in general, which are sorted by eigenvalues in descending order. We applied this approach using functional clustering methods on simulated data and real data to demonstrate the robustness of the efficiency and accuracy of the classification against the MDA clustering algorithms for scRNA-seq data analysis.

## Methods

Our proposed method can be adapted to any existing conventional algorithms such as SC3 [28], Monocle [56, 40, 41], Seurat [55, 8], and SIMLR [59], but does not provide a full workflow/pipeline for scRNA-seq data clustering analysis. After pre-processing and dimensional reduction, based on existing R packages, we build functional data using the index of components and scores from multivariate data in order to increase the classification rates. The phases of analysis in their entirety are schematically illustrated in Figure 1. We reiterate that the goal of this paper is to increase the classification rate by adopting FDA on some analysis phases of conventional scRNA-seq data analysis workflow. The difference is that after dimensional reduction, we build functional data by using a basis function to determine if this change improves the classification rate compared to the results from multivariate data analysis. In figure 1, the analysis phases from A to D are the same processes as the general conventional MDA for scRNA-seq data. Then, we construct functional data after identifying the true dimensionality of a data set (E). Based on converted functional gene expression data, we employ functional clustering algorithms to classify the subpopulation of cell populations using functional cells (F).

**Figure 1:**
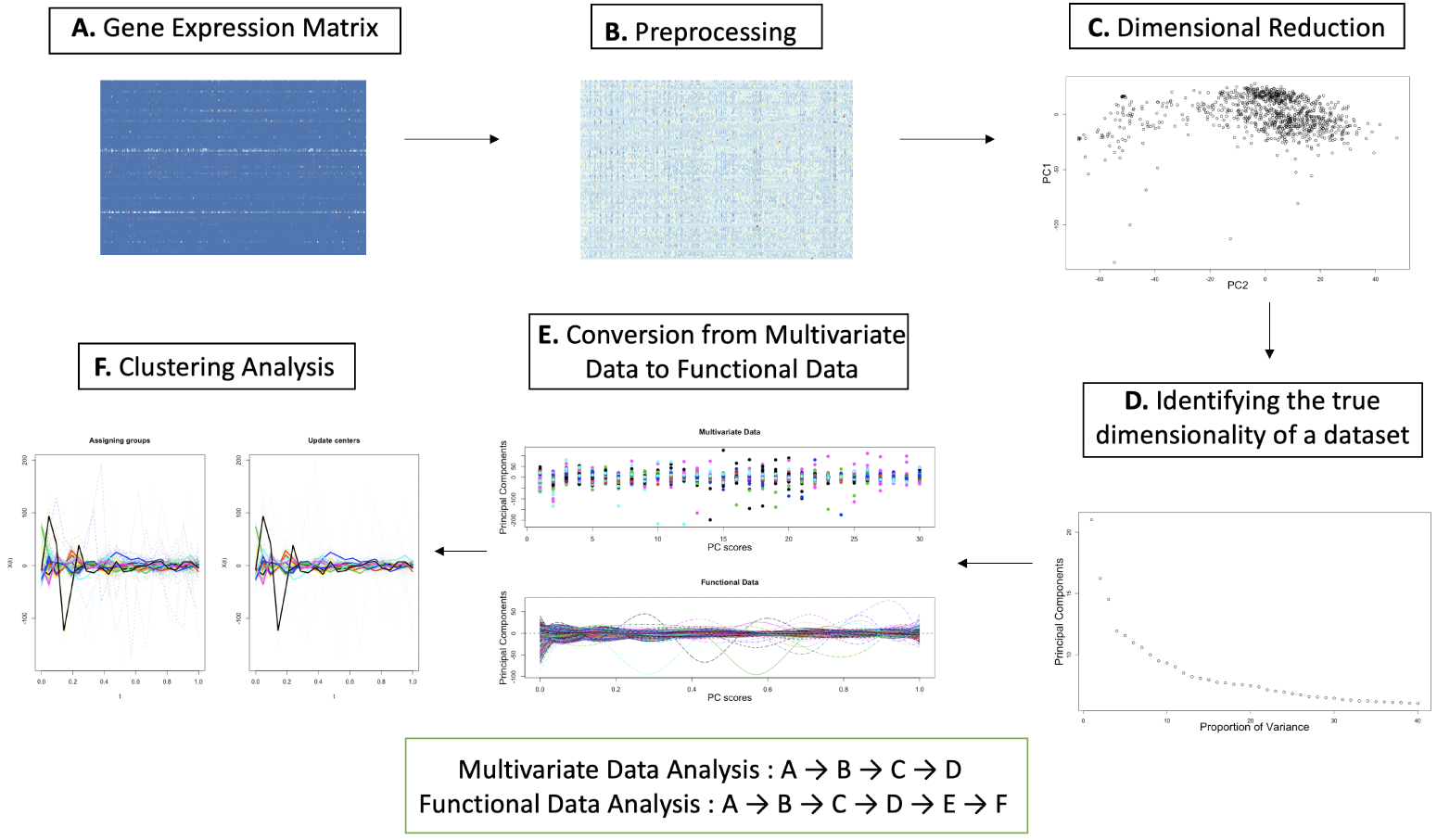
Schematic Overview of the functional data analysis approach. We use existing conventional MDA methods for preprocessing and dimensional reduction. Hence, we adopt existing MDA methods from phases A to D, then based on dimensional reduction results, we convert functional data from multivariate gene expression data.

### Preprocessing Steps

One of the crucial steps in biological experiments, such as scRNA-seq data analysis, is to remove biological or technical errors in the gene expression data [27]. scRNA-seq data analysis is used to explore complex mixtures of cell types in an unsupervised manner. A standard scRNA-seq data analysis involves several tasks that can be performed by various bioinformatics or biostatistics techniques. Zappia et al. [62] categorized these tasks into four broad phases of analysis: data acquisition, data cleaning, cell assignment, and gene identification. The first two phases are generally referred to as the preprocessing steps, and the last two phases are referred to as the statistical analysis steps. Data acquisition can be re-categorized as alignment, de-duplication, and quantification. Data cleaning involves quality control, normalization, and imputation. This work can be done by several existing R packages such as *SC3* [28], *Monocle* [56, 40, 41], and *Seurat* [55, 8]. We implemented these preprocessing steps for scRNA-seq data analysis using the *Seurat* R packages for the downstream analysis. In particular, we normalize and scale the data using *Seurat* for the analysis. Then, we use *scaled data*, which are the z-scored residuals of linear models, to predict gene expression for principal component analysis (PCA) and clustering.

### Framework of building functional data

Functional data analysis was pioneered by Ramsay [42] and then expanded by Ramsay, Silverman, Dalzell, Ferraty and Vieu [47, 44, 46, 14]. A function in functional data analysis is defined in the Hilbert space ℋ, in particular, the 𝕃^2^ space for real square-integrable functions defined on [*a, b*] with the inner product 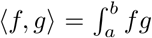. In general, we define a function from the observed multivariate data or functional data with *time* points for the downstream analysis. Then, we apply a smoothing method using a known basis for parametric methods or a kernel function for non-parametric methods. In this paper, using FDA on scRNA-seq data analysis, we no longer consider *time* as a phase of the functions; however, we use the index of the PCs from PCA on the gene expression data.

One of the challenging problems in applying FDA to scRNA-seq data is the presence of high *dropouts*, i.e., zero-inflated counts of gene expression data. This also induces the fitting problem of constructing the function from raw multivariate gene expression data, which produces a tremendous number of spikes when constructing the functional objectives. To solve this issue, we implemented PCA to drastically lower the number of features (dimensions or genes). In this way, we can reduce not only the dimensionality but also the number of dropout values from the scRNA-seq data, which eventually smooth the original data by itself. This is also a common and general step in conventional scRNA-seq data analysis for reducing the dimensionality of the gene expression data. Then, each single cell can now be considered as a single function with PC scores and the index of PCs. The important feature of this analysis is that we treat the index of the PCs as “time points”, i.e., each PC acts analogous to discrete *time* points in the functional gene expression data.

#### Sorting PCs: by eigenvalues

Another problem in functional scRNA-seq data analysis is the order of the PCs, i.e., the phase of the functions, when we build a function from scRNA-seq multivariate data. In MDA, data are considered as discrete vectors; thus, the permutation of the data components is allowed in any statistical analysis. In FDA, however, the permutation of phase components will affect the statistical results in that it should be sorted in order of some characteristics of the data. Traditionally, the most general way to sort the phase components in FDA is in time order, e.g., in seconds, months, or years. For functional scRNA-seq data analysis, we view the PC scores of gene expression profiles as independent realizations of a smooth stochastic process. Hence, the PCs are sorted in descending order according to their variance. Without loss of generality, we discard the number of PCs according to the eigenvalues from the smallest to the largest to reduce the dimensionality of the data. One advantage of using from the largest to smallest eigenvalues here is being able to capture as much of the variance as possible without losing much information from the original data. We consider this framework, i.e., the PCs in descending order, based on the order of the “magnitude” of the corresponding eigenvectors; thus, we treat this strength of variance as the time order to be the functional data. Hence, based on this framework, we build functional data from scRNA-seq data, which is exactly the same as using PCs directly from the results of PCA.

#### Smoothing Functional Data

After building functional gene expression data from discretized multivariate gene expression data, the functional data can be smoothed, leading to the Karhunen-Loéve representation of the observed sample paths as a sum of a smooth mean trend. It can be performed on the functional data, especially where observations are very noisy, to recover the nature of the functional statistics setting. A truncated version of the random part of this representation serves as a statistical approximation of the random process [50]. We first assume that the function *g*(*s*) is observed through the model.

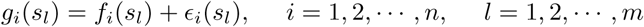

where *ϵ*_*i*_(*s*_*l*_) is the residual error, and *s*_*l*_ is the *l*-th PC, *l* = 1, …, *m*. Then, we can reconstruct the original function *f* (*s*) from the observed function *g*(*s*) using a linear smoother,

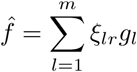

where *ξ*_*lr*_ is the weight that the point *s*_*r*_ gives to the point *s*_*l*_ and *g*_*l*_ = *g*(*s*_*l*_). Then, the function can be smoothed in two ways: using parametric or non-parametric methods.

#### Parametric smoothing method: B-spline basis

A parametric method is also known as a basis representation since we use a known basis to smooth the data. There is no universal basis to use; however, we generally use a B-spline basis and a Fourier basis. A basis is a set of known functions 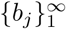 with which any function can be arbitrarily approximated using a linear combination of a sufficiently large number *J* of these functions.

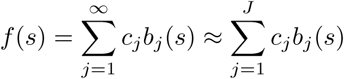

where *c*_*j*_ is the coefficient of the basis function *b*_*j*_.

#### Nonparametric smoothing method: Nadaraya-Watson estimator

Nonparametric smoothing methods, also known as a kernel smoothing method, can be used to represent functional data. In this analysis, we use the Nadaraya-Watson estimator [37] with Gaussian kernel:

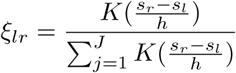

where *K*(·) is the kernel function and *h* is the bandwidth.

#### Generalized Cross-Validation for Smoothing

For parametric and non-parametric smoothing methods, smoothing penalization is crucial for estimating the coefficient of the basis and kernel parameter, respectively. The choice of smoothing parameter is important; however, there is no universal criterion that would ensure an optimal choice. In general, we select the parameter *η* using generalized cross-validation (GCV).

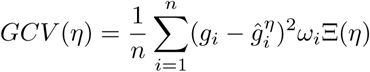

where Ξ(*η*) denotes the type of penalizing function [23] and *ω*_*i*_ is the weight at point *s*_*l*_.

### Clustering Methods

Clustering algorithms are statistical tools used to identify the subpopulation of subjects, such as cell types, in scRNA-seq data analysis. We evaluate three MDA clustering algorithms and three FDA clustering algorithms on gene expression data in this paper. All methods are implemented and publicly available as R packages or scripts. See the references for each method and further details in Table 1.

**Table 1:** Clustering Methods on MDA and FDA.

| Type | Methods | Description | Reference |
| --- | --- | --- | --- |
| MDA | *k*-means | The data given by $x$ are clustered by the *k*-means method, which aims to partition the points into $k$ groups such that the sum of squares from points to the assigned cluster centres is minimized | [25, 15, 31, 33] |
| | hierarchical | A hierarchical cluster analysis using a set of dissimilarities for the $n$ objects being clustered. | [4, 36, 24, 21, 1] |
|  | *mclust* | Model-based clustering based on parameterized finite Gaussian mixture models. | [51, 16, 18, 17] |
| FDA | *k*-means | The method searches the locations around which are grouped data (for a predetermined number of groups) on functional data. | [13, 25] |
|  | *funHDDC* | The *funHDDC* (High-Dimensional Data Clustering) algorithm allows one to cluster functional univariate or multivariate data by modeling each group within a specific functional subspace. | [7] |
|  | *funFEM* | The *funFEM* algorithm allows to cluster functional data by modeling the curves within a common and discriminative functional subspace. | [6] |

For MDA analysis, we perform *k*-means and hierarchical algorithms, which are the most popular clustering algorithms that have been used recently for scRNA-seq data analysis. Many Bioconductors, such as *Seurat, Monocle, and SC3*, perform these clustering techniques to classify the subpopulations to identify and characterize cell populations. In addition to these algorithms, we also apply a recent clustering algorithm, *mclust*, which is a model-based clustering method based on parameterized finite Gaussian mixture models. These models are estimated by the expectation-maximization (EM) algorithm initialized by the hierarchical model-based agglomerative clustering method. Then, we select the optimal model using the Bayesian information criterion (BIC). For clustering methods in FDA, the functional *k*-means method is the same as the one in MDA; however, we define the observations in the Hilbert space, ℋ. *funHDDC* is a model-based algorithm that is based on a functional latent mixture model that fits the functional data in group-specific functional sub-spaces. *funFEM* is also a model-based method but is based on a functional mixture model that allows the clustering of the data in a discriminative functional subspace.

For the stability of the model, we set the random seed to a fixed value internally and externally, and randomly initialize the starting points for optimization 200 times.

## Results

To evaluate the robustness of our approach, we perform the clustering methods that we have described in section 2.3 on both simulated data and real data. For comparison measurement, we calculate the *Hubert-Arabie Adjusted Random Index* (ARI) which is the corrected-for-chance version of the Rand Index [49, 26, 57]:

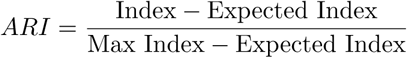

where the independent clusterings have an expected index of zero and identical partitions have an ARI equal to 1. We calculate the ARI using the implementation in the `clues R` package.

### Simulation Study

We generated three simulated data sets using a Gaussian process to observe the improvement in performance of the classification for FDA clustering algorithms. In this simulated experiment, we focused on comparing MDA classification and FDA classification. Therefore, we assume that all the preprocessing and dimensional reduction steps on the scRNA-seq data, such as normalization, scaling, and PCA, are performed. Then, we can only focus on how the FDA approach improves the success rate for classification against MDA classification using PC scores and PCs based on PCA.

We first consider two samples of i.i.d. curves, *X*_*i*_(*s*) and *Y*_*i*_(*s*), generated by independent stochastic processes with different means such that *X*_*i*_(*s*), *Y*_*i*_(*s*) ∈ 𝕃^2^(*I*), where *I* is a compact interval of ℝ. We use Karhunen-Loéve decomposition to generate the sample curves [20, 19, 34]:

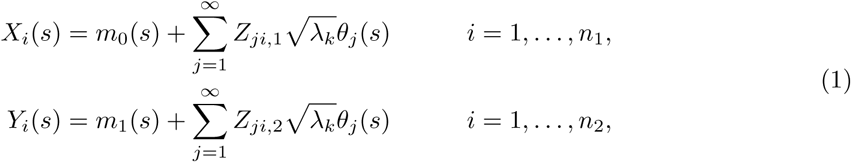

where *s* is the index of PCs (*s*_*l*_, *l* is the *l*-th PC); *m*_0_ and *m*_1_ are the means of each sample for *X*_*i*_ and *Y*_*i*_, respectively; 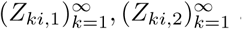 are two sequences of independent standard normal variables; *θ*_*j*_ is the *J/*2 harmonic Fourier basis; λ_*k*_ is the coefficient variance; and *n*_1_ and *n*_2_ are the number of cells in groups 1 and 2, respectively. Because of the infinity of the basis functions, we truncate it into the finite case in terms of *J* known basis functions *θ*_*j*_.

For the initial settings, we generated 150 cell functions for each sample, *X* and *Y*. Hence, we have a total of 300 cells in this simulated data set. Then, we fixed the number of PCs to 40 (*l* = 1, 2, …, 40) assuming that 40 PCs are retained based on some statistical criteria. We set *J* = 40 to have sufficient peaks of the functions for the Fourier basis functions and assign *m*_0_(*s*) = *s*(1 − *s*) for the mean of the sample *X*_*i*_(*s*). For the coefficient variances λ_*k*_, we set

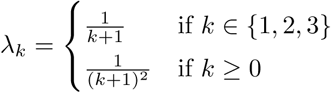

For the mean sample for *Y*_*i*_(*s*), we use three different cases to generate three different data sets to compare the classification efficiency depending on the shape and scale of the functions.

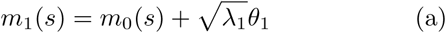

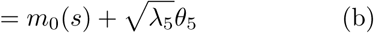

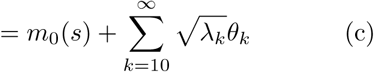

We assigned samples of *X*_*i*_(*s*) and *Y*_*i*_(*s*) as group 1 and group 2, respectively, to group into two different “true” groups. Based on the arguments above, we simulated three different samples of *Y*_*i*_(*s*) for the different means (a), (b), and (c). Figure 2 shows simulated data using Equation 1. The solid red and blue curves are *m*_0_(*s*) and *m*_1_(*s*), which are the means of each sample, respectively. Each panel shows different means of *Y*_*i*_(*s*) for (a), (b), and (c). The difference between the samples in the first panel (case a) is simply the amplitude of the functions. It is easy to see that the shape is very similar and that only the height (*y*-axis) is different. For the second case (case b), we generated a sample of *Y* whereby the mean of its sample lies on the same horizontal line as the mean of sample *X*; however, the shape of the function is different. From the second panel of Figure 2, the shape of the mean of sample *X* is smoother than the shape of the mean of sample *Y* and is a flat curve rather than several distinct peaks in *Y*. The last panel (case c) shows a similar generated function from the second case; however, this time, the sample *Y* has high peaks (spikes) at both the beginning and end of the curve, which give the sample *Y* a very different shape compared to sample *X*. Based on these three different data sets, our goal is to evaluate whether the classification algorithms can cluster into the correct group using the MDA and FDA clustering methods.

**Figure 2:**
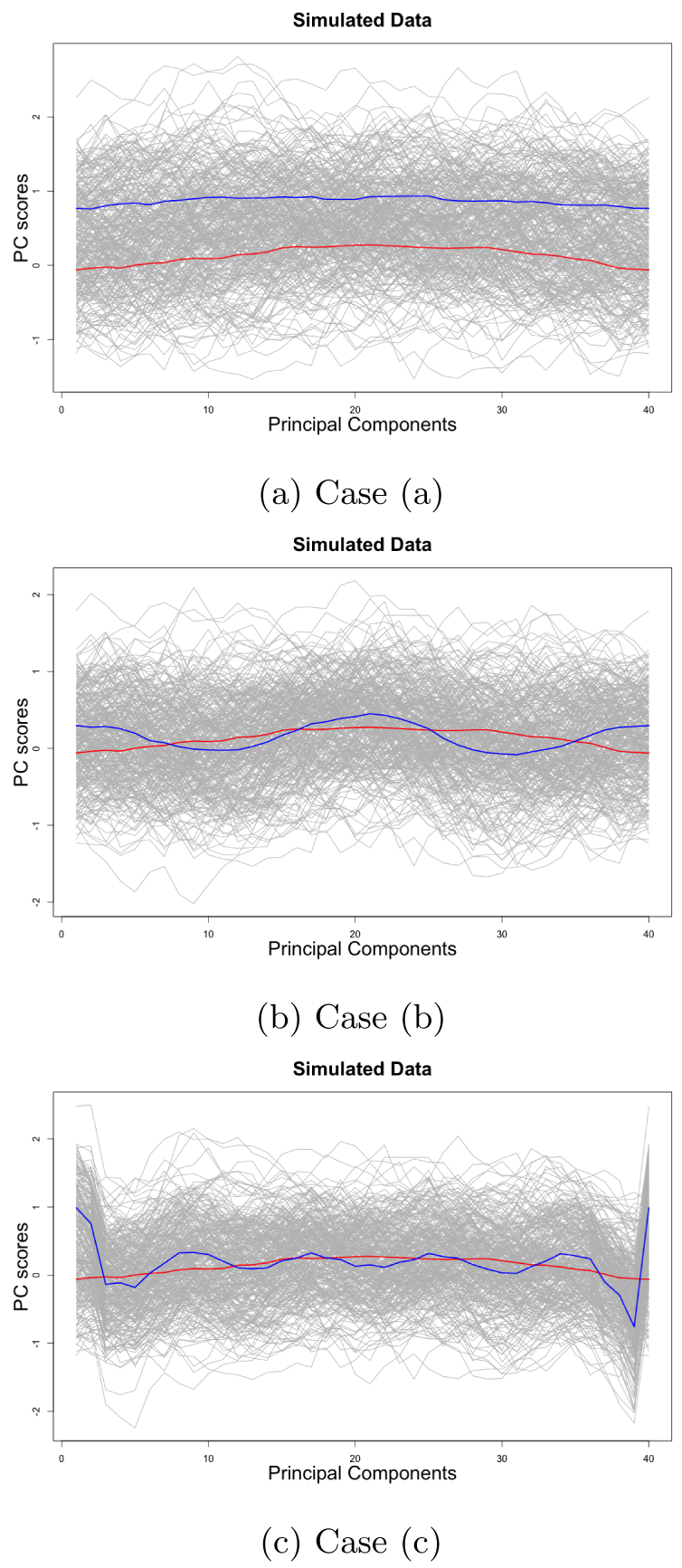
Three simulated data examples with means of *X*_*i*_(*s*) and *Y*_*i*_(*s*) for cases (a), (b), and (c). The gray solid curves are observations. The red curve shows the mean of *X*_*i*_(*s*), and the blue curve shows the mean of *Y*_*i*_(*s*) for each sample. Upon first glance at the observations (gray), the difference between the two samples is not clear due to the noise; however, it is clear that the means for the two samples are different.

Figure 3 shows the observed multivariate data based on generated simulated data, functional data, smoothed functional data using a B-spline basis and a non-parametric method with the Nadaraya-Watson estimator for cases (a), (b), and (c). Here, we use the R packages fda [45, 43, 48] and fda.usc [13] in the R statistical software to convert functional data from the observed multivariate data. Since the observed data have no prominent spikes or outliers, it is difficult to intuitively distinguish between functional data and smoothed functions in these simulated data. However, the number of peaks on both parametric and non-parametric smoothed data is less than functional data without smoothing.

**Figure 3:**
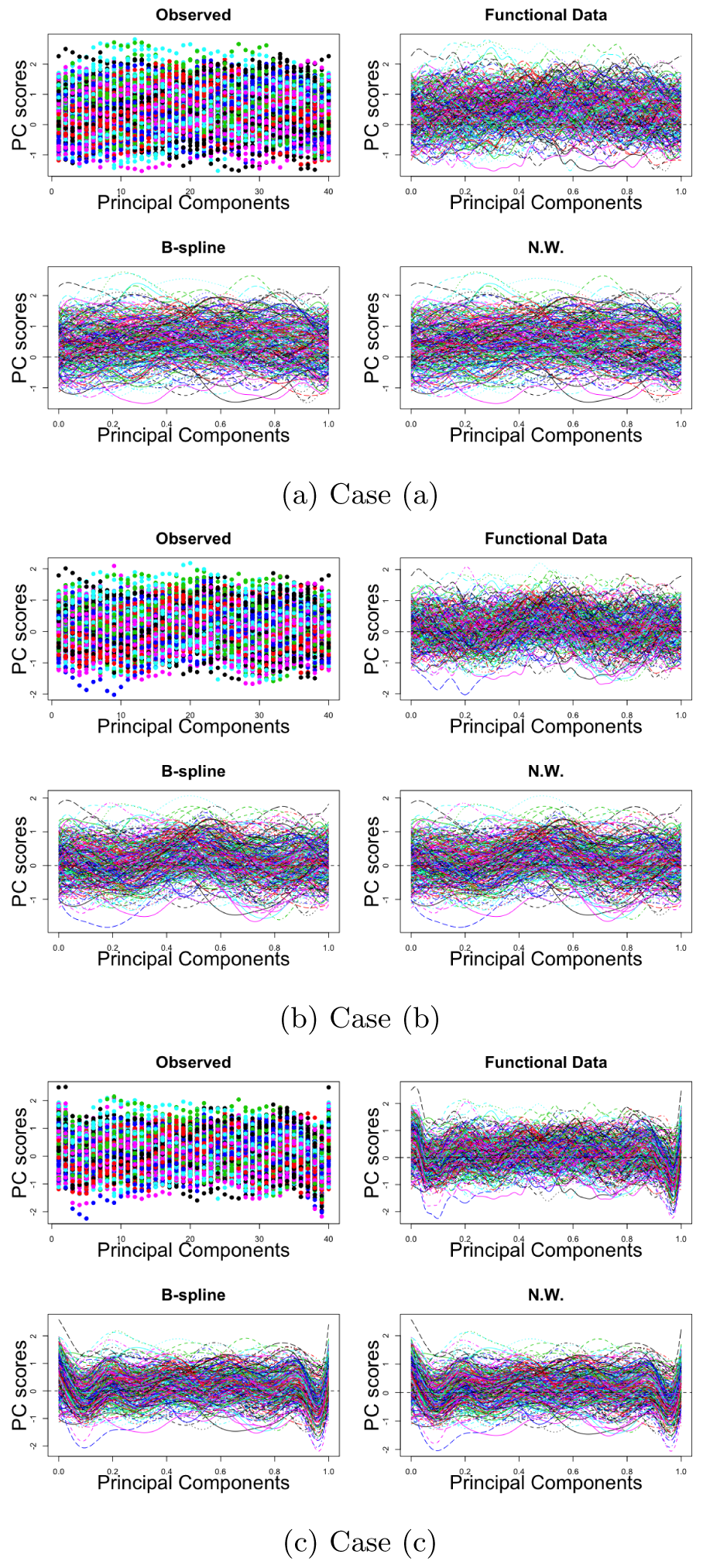
Visualization of each cell with reduced dimensions of simulated gene expression data for cases (a), (b), and (c), respectively. Each color and curve (for functional data) shows the individual cell, where the *x*-axis shows the index of the principal components and the *y*-axis represents the principal component scores. Each panel of the subfigures shows the four observed data sets. The first panel shows the original data, which are considered discrete multivariate data. The second panel is the function data converted from the original discrete data. The third and fourth panels show the smoothed data with B-spline smoothing and kernel smoothing using the Nadaraya-Watson estimator, respectively.

We evaluate the clustering algorithms on these simulated data, and the classification results are shown in Table 2. For MDA, we use three clustering algorithms: the *k*-means, hierarchical, and *mclust* methods. The functional *k*-means, funFEM, and funHDDC clustering algorithms are applied to the functional data. To evaluate the consistency of the classification analysis by switching the phase components of the functions, we randomly sort the order of the components (*unaligned*). Table 2 shows the results of the classification in ARI for each method and each data set. In Table 2, the ARI of applying the functional clustering algorithms outperforms the results of other MDA clustering methods.

**Table 2:** ARI for classification. Bold number shows the highest ARI for each case, (a), (b), and (c), respectively.

| Type | Smoothing | Clustering | (a) |  | (b) |  | (c) |  |
| --- | --- | --- | --- | --- | --- | --- | --- | --- |
|  |  |  | Aligned | Unaligned | Aligned | Unaligned | Aligned | Unaligned |
| FDA | No Particular Smoothing | k-means | 0.282 | 0.358 | -0.003 | -0.000 | -0.002 | 0.008 |
|  |  | funHDDC | 0.479 | 0.048 | 0.018 | -0.002 | 0.054 | 0.020 |
|  |  | funFEM | 0.443 | 0.382 | 0.010 | 0.000 | 0.054 | 0.013 |
|  | Parametric (b-spline) | k-means | 0.408 | 0.391 | -0.001 | -0.002 | 0.007 | -0.002 |
|  |  | funHDDC | **0.488** | 0.434 | **0.034** | 0.004 | **0.934** | 0.008 |
|  |  | funFEM | 0.479 | 0.425 | 0.032 | -0.002 | 0.883 | 0.001 |
|  | Non-Parametric (NW) | k-means | 0.350 | 0.304 | 0.000 | -0.001 | 0.005 | 0.000 |
|  |  | funHDDC | **0.488** | 0.443 | **0.034** | -0.001 | 0.203 | 0.008 |
|  |  | funFEM | 0.479 | 0.434 | 0.032 | 0.010 | 0.197 | -0.003 |
| MDA |  | k-means | 0.434 |  | 0.004 |  | 0.011 |  |
|  |  | hierarchical | 0.311 |  | 0.003 |  | -0.003 |  |
|  |  | mclust | 0.434 |  | 0.004 |  | -0.003 |  |

In the comparison of the three cases (a), (b), and (c), case (c) shows the highest classification rate (0.934) since the shape and peaks or valleys on both sides affect the differences between the samples *X* and *Y*. Case (a) shows the second-highest ARI (0.488) among the three data sets. This implies that the height (or *y*-axis or vertical difference) also plays a major role in grouping the observations into the correct group. Case (b) shows the lowest accuracy (0.034) due to the similarity between the samples *X* and *Y*. As expected, the average of the classification rates for the unaligned phase components of the functions is lower than the results of the aligned functional data. Particularly, case (c) shows the large difference in the ARI between aligned and unaligned functional data. This implies that the order of the phase components of the function is also a major factor in applying functional data analysis methods to achieve the highest accuracy rate.

### Application to Real Data

The real scRNA-seq data sets were collected from *conquer* [54] and used for our classification evaluations: GSE 66507 (here denoted **Blakeley2015**) [5], GSE 52529-GPL16791 (**Trapnell**) [56], GSE 74596 (**Engel2016**) [12], GSE 66053-GPL 18573 (**PadovanMerhar2015**) [39], GSE 62270-GPL17021 (**Grun2015**) [22], GSE 81903 (**Shekhar2016**) [53], GSE 102299 (**Wallrapp2017**) [58], and GSE 48968-GPL 17021-125bp (**Shalek2014**) [52]. These data sets are not expected to be used with the aim of detecting subpopulations of cell populations. Hence, the cluster labels known as “true” cluster labels might not represent the strongest signal present in the data [11]. In other words, general classification algorithms cannot detect the transcriptional signal or the characteristics of each cluster, which statistically derives different cluster labels rather than true cluster labels. Duó et al. [11] noted that these labels can be biased in favor of current methodologies. Therefore, our goal for this real data analysis is to detect the *true* subpopulations. Hence, it is important to validate the performance of functional clustering algorithms considering each cell as a functional shape using these data sets to uncover the functional nature of scRNA-seq gene expression data.

The descriptions of each data set, including the number of cells and subpopulations, are shown in Table 3. For example, **Blakeley2016, Trapnell2014, Engel2016, PadovanMerhar2015, Grun2015, Shekhar2016, Wallrapp2017** and **Shalek2014** have 3, 3, 4, 4, 4, 4, 3, and 6 subpopulations, respectively. The selected cell phenotype was used to define the “true” partition of cells when evaluating the clustering methods. The details of the subpopulations and the methods for finding these true subpopulations for each data set are explained and described in [54].

**Table 3:** Description of scRNA-seq data.

| Data set | Organism | Sequencing Protocol | Cells | Subpopulations | Description | References |
| --- | --- | --- | --- | --- | --- | --- |
| **Blakeley2015** | Homo Sapiens | SMARTer | 30 | 3 | Preimplantation embryos | [5] |
| **Trapnell2014** | Homo Sapiens | SMARTer C1 | 288 | 3 | Primary myoblasts over a time course of serum-induced differentiation | [56] |
| **Engel2016** | Mus Musculus | Smart-Seq2 | 203 | 4 | purified populations of thymic NKT cell subsets | [12] |
| **PadovanMerhar2015** | Homo Sapiens | Fluidigm C1 Auto Prep | 96 | 4 | Live (large and medium) and fixed single cell | [39] |
| **Grun2015** | Mus Musculus | CEL-Seq | 2891 | 4 | Cells from mouse intestinal organoids | [22] |
| **Shekhar2016** | Mus Musculus | Smart-Seq2 | 384 | 4 | P17 retinal cells from the Kcng4-cre;stop-YFP X Thy1-stop-YFP Line#1 mice | [53] |
| **Wallrapp2017** | Mus Musculus | Smart-Seq2 | 752 | 3 | innate lymphoid cells from mouse lungs after various treatments | [58] |
| **Shalek2014** | Mus Musculus | SMARTer C1 | 935 | 6 | Dendritic cells stimulated with pathogenic components | [52] |

For the preprocessing steps, such as quality control, normalization, and scaling, we use the *Seurat* R package [55, 8] to perform the downstream analysis. *Seurat* can also perform *t-SNE* analysis and clustering methods, such as *k*-means; however, we did not implement these clustering methods using *Seurat* and rather used general-purpose R packages or scripts (Table 1) for clustering the cell subpopulations. More details about these preprocessing steps and procedures using the *Seurat* Bioconductor are described in [8].

After applying the preprocessing steps on the scRNA-seq data, we used all genes to perform PCA to reduce the dimensionality of the gene expression data. In this analysis, we can visualize the distribution or pattern of the cell populations by plotting PC1 vs. PC2. Figure 4 shows the PC1 vs. PC2 plot for each scRNA-seq data set after performing PCA. In this figure, it is complicated to group a set of objects into the “true” groups as given in Table 3 without any statistical clustering analysis. One of the main reasons for these results is that the “true” cluster labels do not represent the strongest signal present in the multivariate data. For example, **Blakeley2015, Trapnell2014, Engel2016, PadovanMerhar2015, Grun2015, Shekhar2016, Wallrapp2017**, and **Shalek2014** data sets have 3, 3, 4, 4, 4, 4, 3, and 6 subpopulations, respectively; however, none of the plots in Figure 4 show a clear distinctive number of clusters for each data set. In this sense, we want to see how FDA, which considers each cell as a functional shape, performs in classification against multivariate data for these phenomena whereby the signal of the “true” labels is not sufficiently strong. This functional approach will identify the hidden functional nature of scRNA-seq data.

**Figure 4:**
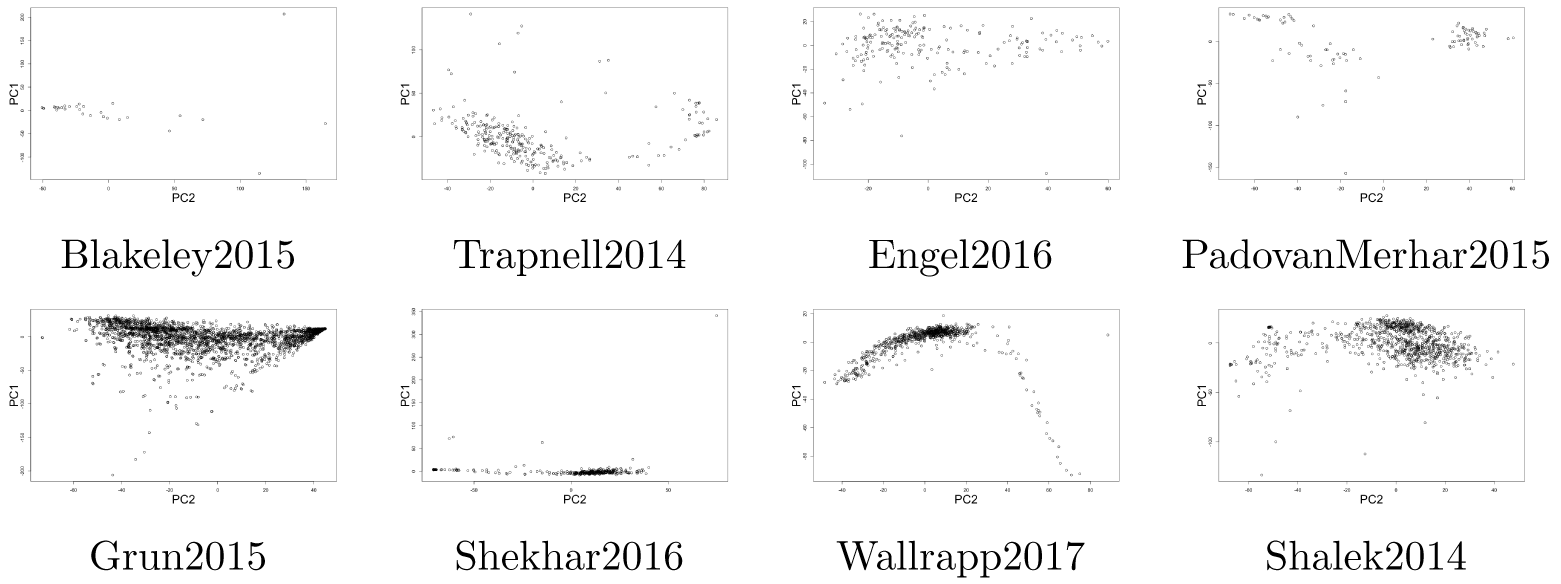
PC1 vs. PC2 plot after performing Principal Component Analysis for each scRNA-seq data set. None of the data sets could be separated into “true” numbers of subpopulations from the PC 1 vs. PC2 plot. None of the plots in Figure 4 show a clear distinctive number of clusters for each data set. In this sense, we want to see how FDA, which considers each cell as a functional shape, performs in classification against multivariate data for these phenomena whereby the signal of the “true” labels is not sufficiently strong. This functional approach will identify the hidden functional nature of scRNA-seq data.

We visualize the scree plot and perform a jackstraw [9] to determine the optimal number of PCs to reduce the dimensionality of the original data. A scree plot in PCA is a useful tool that visualizes saturation in the relationship between the number of PCs and the percentage of the variance explained. We generally decide the number of principal components that corresponds to the “elbow” part of the curve to have sufficient information of the original data. In this experiment, we chose from PC1 to PC20 -PC40 for the real data sets for the downstream analysis.

One of the important features of the function in the Hilbert space, ℋ, is that unlike the vectors in MDA, it does not allow the permutations of the components, i.e., the phase components. Therefore, we perform and build the function where phase components are sorted according to the eigenvalues. It is the simplest way that we can directly use PCs from PCA results. Then scRNA-seq gene expression data are reconstructed after PCA, where the *x*-axis represents the PCs while the *y*-axis represents the PC scores for each cell. Then we build the functional data using `fda` and `fda.usc` in the R software to convert the functional data from the original data. Then, we apply two functional smoothing methods, parametric and nonparametric, to smooth the functional data. We set a sufficient number of bases to construct the functional data from the scRNA-seq data.

We performed clustering analysis on real data. The classification results and accuracy rates ARI are shown in Table 4. In these real data experiments, we utilized two different cases of phase alignment for each data set: 1) randomly aligned (*Unaligned*) and 2) aligned (*Aligned*). In this table, the bold number represents the highest accuracy (ARI) for each scRNA-seq data set.

**Table 4:** Classification results for real data. Each bolded number shows the highest ARI for each data set.

|  | Type | FDA |  |  |  |  |  |  |  |  | MDA |  |  |
| --- | --- | --- | --- | --- | --- | --- | --- | --- | --- | --- | --- | --- | --- |
|  | Smoothing | No Smoothing |  |  | Parametric (B-Spline) |  |  | Non-parametric (N.W.) |  |  |  |  |  |
|  | Clustering | funHDDC | k-means | funFEM | funHDDC | k-means | funFEM | funHDDC | k-means | funFEM | k-means | hierarchical | mclust |
| Blakeley2015 | Aligned | -0.034 | -0.023 | **0.119** | 0.009 | -0.089 | -0.027 | -0.036 | -0.007 | -0.036 | -0.078 | -0.059 | -0.038 |
|  | Unaligned | -0.065 | -0.059 | 0.045 | -0.024 | -0.037 | 0.005 | -0.004 | -0.007 | 0.023 |  |  |  |
| Trapnell2014 | Aligned | **0.227** | 0.014 | -0.001 | -0.000 | -0.000 | 0.023 | 0.027 | 0.005 | 0.005 | -0.001 | 0.000 | 0.040 |
|  | Unaligned | 0.041 | -0.001 | -0.000 | 0.126 | 0.003 | 0.118 | 0.219 | 0.005 | 0.002 |  |  |  |
| Engel2016 | Aligned | **0.648** | 0.039 | 0.344 | 0.042 | 0.018 | 0.062 | 0.071 | 0.065 | 0.047 | 0.308 | 0.001 | 0.477 |
|  | Unaligned | 0.459 | 0.039 | 0.198 | 0.291 | 0.018 | 0.174 | 0.124 | 0.064 | 0.270 |  |  |  |
| PadovanMerhar2015 | Aligned | -0.074 | -0.047 | **0.242** | 0.104 | 0.202 | 0.156 | 0.001 | -0.069 | 0.202 | 0.079 | -0.047 | 0.136 |
|  | Unaligned | 0.106 | -0.047 | 0.133 | 0.091 | 0.202 | 0.163 | 0.121 | -0.095 | 0.227 |  |  |  |
| Grun2015 | Aligned | 0.190 | 0.087 | **0.223** | 0.057 | 0.055 | 0.134 | 0.075 | 0.046 | 0.134 | 0.172 | -0.000 | 0.122 |
|  | Unaligned | 0.162 | 0.110 | 0.147 | 0.105 | 0.070 | 0.119 | 0.078 | 0.020 | 0.117 |  |  |  |
| Shekhar2016 | Aligned | **0.221** | 0.008 | 0.015 | 0.019 | 0.113 | 0.048 | 0.019 | 0.006 | 0.048 | 0.008 | -0.000 | 0.011 |
|  | Unaligned | 0.116 | 0.008 | 0.012 | 0.145 | 0.007 | 0.107 | 0.143 | 0.021 | 0.102 |  |  |  |
| Wallrapp2017 | Aligned | 0.024 | 0.052 | 0.184 | **0.320** | 0.064 | 0.135 | **0.320** | 0.034 | 0.139 | 0.238 | 0.000 | 0.040 |
|  | Unaligned | 0.052 | 0.286 | 0.138 | -0.001 | 0.064 | 0.213 | 0.051 | 0.034 | 0.297 |  |  |  |
| Shalek2014 | Aligned | 0.203 | 0.028 | 0.160 | 0.125 | 0.178 | **0.307** | 0.078 | 0.116 | 0.150 | 0.222 | -0.001 | 0.102 |
|  | Unaligned | 0.157 | 0.226 | 0.171 | 0.183 | 0.178 | 0.249 | 0.123 | 0.084 | 0.115 |  |  |  |

#### Functional data performed better than multivariate data

Our main goal in this paper is to see the improvement of the classification rate by considering a cell as a single-entity, functional structure rather than discrete multivariate data. As expected, the functional data approach outperformed in classification compared to the clustering results from MDA. In Table 4, all of the data sets show higher accuracy rates for functional clustering algorithms, mostly when functions are not smoothed. In particular, the first six data sets show the highest ARI when functions are not smoothed in Table 4. The last two data sets, **Wallrapp2017** and **Shalek2014**, have the highest ARI when the functions are smoothed. For **Wall-rapp2017** data, both parametric and non-parametric show the same classification results. These results indicate that the first six data sets might have small biological or technical noises where the smoothing method actually removed some structure from the data that might be important to the analysis. This might be because the gene expression data are already normalized and scaled in order to remove some noises in preprocessing step. As a result, the smoothed functional data classification analyses cannot achieve the highest accuracy performance compared to the ones without smoothing.

#### Functional data sorted by eigenvalues showed the highest ARI

We are also interested in the improvement of classification rates by ordering the PCs for the phase components. We proposed the criterion that in order to achieve the best performance, sorting the PCs by eigenvalues would have the highest accuracy in classification. Table 4 shows that all of the data sets achieved the highest ARI when the PCs are sorted by eigenvalues rather than randomly, which means that our proposed criterion obtains the best classification rate when we apply the functional clustering algorithm to functional data converted from multivariate gene expression data.

## Discussion

In this study, we have proposed a new framework, functional data analysis for scRNA-seq data, to identify the subpopulations of cell populations. We have demonstrated that functional data analysis can be applied to scRNA-seq data to improve the accuracy rate of classification to identify and characterize cell populations. We consider a cell as a functional shape rather than discrete vector. This framework improves the classification rates in scRNA-seq data analysis, in particular, when the biological data may not represent the strongest signal present in the data. This was one of the major problems in MDA since any evaluation is based on typical inference by clustering the cells using MDA clustering algorithms in which clustering labels can be biased [11]. In MDA, most bioinformatics techniques and methods are focused on examining the discrete genomic units of genes, and this approach might ignore the functional nature of the data and eventually loses important information such as the dependency between the phase components that reflects the gene expression data. In this sense, an approach based on FDA also plays a major role in scRNA-seq data analysis.

This novel proposed approach is to apply functional data analysis when multivariate data are very noisy and complex. Generally, we convert from multivariate data to functional data and then perform functional principal component analysis (fPCA) to reduce the dimension of the functional data. However, problems arise when the raw multivariate data are too complex. In our approach, we consider how to handle the number of spikes that are considered as dropouts for gene expression data in scRNA-seq data analysis. To solve this problem, we first perform PCA in MDA to condense the gene expression data for formatting for functional data. In this way, we can not only reduce the dimensionality of the data but also remove the dropouts such that the function can be easily fitted to the data.

There are some limitations in our functional scRNA-seq data analysis. Due to the removal of the noise several times, some of the important information may be ignored in the analysis. For example, we first normalize, scale, and filter the cells in gene expression data to reduce biological errors, such as batch effects; then, we perform PCA to remove the dropouts of the data to fit the function. Then, we fit the function on discretized gene expression data using a known basis. We also apply smoothing techniques to smooth the functional data. When implementing these several steps, we might remove crucial information about the gene expression data. This indicates the need to exercise careful caution when using smoothing methods for functional gene expression data.

## Conclusions

This new statistical technique enhances the classification performance and ultimately improves the understanding of stochastic biological processes. Therefore, this new framework provides an essentially different scRNA-seq data analytical approach, which can complement conventional MDA methods such as Seurat, SC3, and Monocle. It can be truly effective when current MDA methods cannot detect or uncover the hidden nature of the gene expression dynamics due to weak signals by adapting these methods and constructing functional data after dimensional reduction. Moreover, this study enables the conversion of functional data from multivariate gene expression data, and any further functional statistical analysis is applicable to scRNA-seq data analysis. This is a critical step for scRNA-seq data analysis as well as functional data analysis.

## Ethics approval and consent to participate

Not Applicable

## Consent for publication

Not Applicable

## Availability of data and materials

The scRNA-seq data sets are publicly available from *Conquer*: http://imlspenticton.uzh.ch:3838/conquer/. The `R` software is available from https://www.r-project.org/ and the `R` packages are available from https://cran.r-project.org/.

## Author’s contributions

KA and HF initiated the study. KA designed and implemented R code. KA and HF analyzed and interpreted the results. KA wrote the original draft and HF reviewed and edited the manuscript.

## Acknowledgements

We would like to thank the members of the Fujiwara laboratory for valuable discussion.

## References

1. M. R. Anderberg. Cluster analysis for applications: probability and mathematical statistics: a series of monographs and textbooks, volume 19. Academic press, 2014.

2. T. S. Andrews and M. Hemberg. Identifying cell populations with scrnaseq. Molecular aspects of medicine, 59:114–122, 2018.

3. Z. Bar-Joseph, G. K. Gerber, D. K. Gifford, T. S. Jaakkola, and I. Simon. Continuous representations of time-series gene expression data. Journal of Computational Biology, 10(3-4):341–356, 2003.

4. R. Becker. The new S language. CRC Press, 2018.

5. P. Blakeley, N. Fogarty, I. Del Valle, S. E. Wamaitha, T. X. Hu, K. Elder, P. Snell, L. Christie, P. Robson, and K. K. Niakan. Defining the three cell lineages of the human blastocyst by single-cell rna-seq. Development, 142(18):3151–3165, 2015.

6. C. Bouveyron, E. Côme, J. Jacques, et al. The discriminative functional mixture model for a comparative analysis of bike sharing systems. The Annals of Applied Statistics, 9(4):1726–1760, 2015.

7. C. Bouveyron and J. Jacques. Model-based clustering of time series in group-specific functional subspaces. Advances in Data Analysis and Classification, 5(4):281–300, 2011.

8. A. Butler, P. Hoffman, P. Smibert, E. Papalexi, and R. Satija. Integrating single-cell transcriptomic data across different conditions, technologies, and species. Nature biotechnology, 36(5):411, 2018.

9. N. C. Chung and J. D. Storey. Statistical significance of variables driving systematic variation in high-dimensional data. Bioinformatics, 31(4):545–554, 2014.

10. N. Coffey, J. Hinde, and E. Holian. Clustering longitudinal profiles using p-splines and mixed effects models applied to time-course gene expression data. Computational Statistics & Data Analysis, 71:14–29, 2014.

11. A. Duo, M. D. Robinson, and C. Soneson. A systematic performance evaluation of clustering methods for single-cell rna-seq data. F1000Research, 7, 2018.

12. I. Engel, G. Seumois, L. Chavez, D. Samaniego-Castruita, B. White, A. Chawla, D. Mock, P. Vijayanand, and M. Kronenberg. Innate-like functions of natural killer t cell subsets result from highly divergent gene programs. Nature immunology, 17(6):728, 2016.

13. M. Febrero-Bande and M. Oviedo de la Fuente. Statistical computing in functional data analysis: the r package fda. usc. Journal of Statistical Software, 51(4):1–28, 2012.

14. F. Ferraty and P. Vieu. Nonparametric functional data analysis: theory and practice. Springer Science & Business Media, 2006.

15. E. Forgey. Cluster analysis of multivariate data: Efficiency vs. interpretability of classification. Biometrics, 21(3):768–769, 1965.

16. C. Fraley and A. E. Raftery. Model-based clustering, discriminant analysis, and density estimation. Journal of the American statistical Association, 97(458):611–631, 2002.

17. C. Fraley and A. E. Raftery. Bayesian regularization for normal mixture estimation and model-based clustering. Journal of classification, 24(2):155–181, 2007.

18. C. Fraley, A. E. Raftery, T. B. Murphy, and L. Scrucca. mclust version 4 for r: normal mixture modeling for model-based clustering, classification, and density estimation. Technical report, Technical report, 2012.

19. A. Ghiglietti, F. Ieva, and A. M. Paganoni. Statistical inference for stochastic processes: two-sample hypothesis tests. Journal of Statistical Planning and Inference, 180:49–68, 2017.

20. A. Ghiglietti and A. M. Paganoni. Statistical inference for functional data based on a generalization of mahalanobis distance. Mox Report 39/2014, Department of Mathematics, Politecnico di Milano, 6, 2014.

21. A. D. Gordon. A review of hierarchical classification. Journal of the Royal Statistical Society: Series A (General), 150(2):119–137, 1987.

22. D. Grün, A. Lyubimova, L. Kester, K. Wiebrands, O. Basak, N. Sasaki, H. Clevers, and A. van Oudenaarden. Single-cell mrna sequencing reveals rare intestinal cell types. ncbi geo database. 2015.

23. W. Härdle. Applied nonparametric regression. Number 19. Cambridge university press, 1990.

24. J. A. Hartigan. Clustering algorithms. 1975.

25. J. A. Hartigan and M. A. Wong. Algorithm as 136: A k-means clustering algorithm. Journal of the Royal Statistical Society. Series C (Applied Statistics), 28(1):100–108, 1979.

26. L. Hubert and P. Arabie. Comparing partitions. Journal of classification, 2(1):193–218, 1985.

27. B. Hwang, J. H. Lee, and D. Bang. Single-cell rna sequencing technologies and bioinformatics pipelines. Experimental & molecular medicine, 50(8):96, 2018.

28. V. Y. Kiselev, K. Kirschner, M. T. Schaub, T. Andrews, A. Yiu, T. Chandra, K. N. Natarajan, W. Reik, M. Barahona, A. R. Green, et al. Sc3: consensus clustering of single-cell rna-seq data. Nature methods, 14(5):483, 2017.

29. R. R. Klevecz and D. B. Murray. Genome wide oscillations in expression–wavelet analysis of time series data from yeast expression arrays uncovers the dynamic architecture of phenotype. Molecular Biology Reports, 28(2):73–82, 2001.

30. X. Leng and H-G Müller. Classification using functional data analysis for temporal gene expression data. Bioinformatics, 22(1):68–76, 2005.

31. S. Lloyd. Least squares quantization in pcm. IEEE transactions on information theory, 28(2):129–137, 1982.

32. Y. Luan and H. Li. Clustering of time-course gene expression data using a mixed-effects model with b-splines. Bioinformatics, 19(4):474–482, 2003.

33. J. MacQueen et al. Some methods for classification and analysis of multivariate observations. In Proceedings of the fifth Berkeley symposium on mathematical statistics and probability, volume 1, pages 281–297. Oakland, CA, USA, 1967.

34. A. Martino. Classification algorithms for multivariate functional data. 2016.

35. V. Menon. Clustering single cells: a review of approaches on high-and low-depth single-cell rna-seq data. Briefings in functional genomics, 17(4):240–245, 2017.

36. F. Murtagh and P. Legendre. Ward’s hierarchical agglomerative clustering method: which algorithms implement ward’s criterion? Journal of classification, 31(3):274–295, 2014.

37. E. A. Nadaraya. On estimating regression. Theory of Probability & Its Applications, 9(1):141–142, 1964.

38. R. Opgen-Rhein and K. Strimmer. Learning causal networks from systems biology time course data: an effective model selection procedure for the vector autoregressive process. BMC bioinformatics, 8(2):S3, 2007.

39. O. Padovan-Merhar, G. P. Nair, A. G. Biaesch, A. Mayer, S. Scarfone, S. W. Foley, A. R. Wu, L. S. Churchman, A. Singh, and A. Raj. Single mammalian cells compensate for differences in cellular volume and dna copy number through independent global transcriptional mechanisms. Molecular cell, 58(2):339–352, 2015.

40. X. Qiu, A. Hill, J. Packer, D. Lin, Y-A Ma, and C. Trapnell. Single-cell mrna quantification and differential analysis with census. Nature methods, 14(3):309, 2017.

41. X. Qiu, Q. Mao, Y. Tang, L. Wang, R. Chawla, H. A. Pliner, and C. Trapnell. Reversed graph embedding resolves complex single-cell trajectories. Nature methods, 14(10):979, 2017.

42. J. O. Ramsay. When the data are functions. Psychometrika, 47(4):379–396, 1982.

43. J. O. Ramsay. Functional data analysis. Encyclopedia of Statistics in Behavioral Science, 2005.

44. J. O. Ramsay and C. J. Dalzell. Some tools for functional data analysis. Journal of the Royal Statistical Society. Series B (Methodological), 53(3):539–572, 1991.

45. J. O. Ramsay, G. Hooker, and S. Graves. Functional data analysis with r and matlab, vol. 66, 2010.

46. J. O. Ramsay and B. W. Silverman. Applied functional data analysis: methods and case studies, volume 77. Citeseer, 2002.

47. J. O. Ramsay and B. W. Silverman. Functional Data Analysis. Springer, 2nd edition, 2005.

48. J. O. Ramsay and B. W. Silverman. Applied functional data analysis: methods and case studies. Springer, 2007.

49. W. M. Rand. Objective criteria for the evaluation of clustering methods. Journal of the American Statistical association, 66(336):846–850, 1971.

50. J. A. Rice and B. W. Silverman. Estimating the mean and covariance structure nonparametrically when the data are curves. Journal of the Royal Statistical Society: Series B (Methodological), 53(1):233–243, 1991.

51. L. Scrucca, M. Fop, T. B. Murphy, and A. E. Raftery. mclust 5: clustering, classification and density estimation using gaussian finite mixture models. The R journal, 8(1):289, 2016.

52. A. K. Shalek, R. Satija, J. Shuga, J. J. Trombetta, D. Gennert, D. Lu, P. Chen, R. S. Gertner, J. T. Gaublomme, N. Yosef, et al. Single-cell rna-seq reveals dynamic paracrine control of cellular variation. Nature, 510(7505):363, 2014.

53. K. Shekhar, S. W. Lapan, I. E. Whitney, N. M. Tran, E. Z. Macosko, M. Kowalczyk, X. Adiconis, J. Z. Levin, J. Nemesh, M. Goldman, et al. Comprehensive classification of retinal bipolar neurons by single-cell transcriptomics. Cell, 166(5):1308–1323, 2016.

54. C. Soneson and M. D. Robinson. Bias, robustness and scalability in single-cell differential expression analysis. Nature methods, 15(4):255, 2018.

55. T. Stuart, A. Butler, P. Hoffman, C. Hafemeister, E. Papalexi, W. M. Mauck III, Y. Hao, M. Stoeckius, P. Smibert, and R. Satija. Comprehensive integration of single-cell data. Cell, 177(7):1888–1902, 2019.

56. C. Trapnell, D. Cacchiarelli, J. Grimsby, P. Pokharel, S. Li, M. Morse, N. J. Lennon, K. J. Livak, T. S. Mikkelsen, and J. L. Rinn. The dynamics and regulators of cell fate decisions are revealed by pseudotemporal ordering of single cells. Nature biotechnology, 32(4):381, 2014.

57. N. X. Vinh, J. Epps, and J. Bailey. Information theoretic measures for clusterings comparison: is a correction for chance necessary? In Proceedings of the 26th annual international conference on machine learning, pages 1073–1080. ACM, 2009.

58. A. Wallrapp, S. J. Riesenfeld, P. R. Burkett, E. Abdulnour, J. Nyman, D. Dionne, M. Hofree, M. S. Cuoco, C. Rodman, and D. and others Farouq. The neuropeptide nmu amplifies ilc2-driven allergic lung inflammation. Nature, 549(7672):351, 2017.

59. B. Wang, J. Zhu, E. Pierson, D. Ramazzotti, and S. Batzoglou. Visualization and analysis of single-cell rna-seq data by kernel-based similarity learning. Nature methods, 14(4):414, 2017.

60. S. Wesolowski, D. Vera, and W. Wu. Srsf shape analysis for sequencing data reveal new differentiating patterns. Computational biology and chemistry, 70:56–64, 2017.

61. P-S Wu and H-G Müller. Functional embedding for the classification of gene expression profiles. Bioinformatics, 26(4):509–517, 2010.

62. L. Zappia, B. Phipson, and A. Oshlack. Exploring the single-cell rna-seq analysis landscape with the scrna-tools database. PLoS computational biology, 14(6):e1006245, 2018.

